# Multimodal characterization of transcriptionally defined ventral tegmental area dopamine neurons

**DOI:** 10.64898/2026.03.16.712214

**Authors:** N. Dalton Fitzgerald, Emily T. Jorgensen, Catherine E. Newman, Lydia G. Slocum, Katelyn M. Varden, Jeremy J. Day

**Affiliations:** Department of Neurobiology, University of Alabama at Birmingham, Birmingham, AL 35294, USA

## Abstract

Ventral tegmental area (VTA) dopamine (DA) neurons are highly implicated in reward learning, motivated behaviors, and substance use disorders. DA neurons in the VTA are traditionally characterized by expression of genes involved in DA synthesis, release, or reuptake, such as tyrosine hydroxylase (encoded by the *Th* gene), which is the rate-limiting step in DA synthesis. However, recent transcriptomic studies have revealed substantial cellular heterogeneity within the VTA, including multiple subtypes of VTA DA neurons. Using single nucleus RNA sequencing, we previously identified two transcriptionally distinct *Th*+ subpopulations: a DA/glutamate/GABA Combinatorial neuron marked by *Slc26a7* and a DA-only neuron marked by *Gch1*. However, the functional properties of these distinct DA neuron classes remain unknown. Here, we developed an AAV-based strategy enabling cell-type-specific access to these populations and performed comparative transcriptional, electrophysiological, and anatomical analyses, providing the first functional characterization of these transcriptionally-defined DA neuron subtypes. Whole-cell recordings revealed similar baseline membrane properties but a divergence in intrinsic excitability and latency to fire action potentials after current input. Anatomical mapping revealed overlapping but biased projection patterns, and Combinatorial neurons, but not DA-only neurons, were selectively recruited following experience with cocaine. Together, these findings reveal functional specialization among transcriptionally-defined VTA DA neuron subpopulations, dissociating DA-specific from multi-neurotransmitter properties and refining our understanding of VTA heterogeneity.

## INTRODUCTION

The ventral tegmental area (VTA) is a midbrain region critical for motivation, reward processing, and reinforcement learning. Dysregulation of the VTA has been implicated in many psychiatric diseases, such as substance use disorders (SUD), schizophrenia, and depression, accentuating the need to further understand this region. While the role of the VTA in motivation has historically been attributed to dopamine (DA), recent studies have highlighted vast heterogeneity within this midbrain region. DA neurons are classically identified by expression of tyrosine hydroxylase (TH), the dopamine transporter (DAT), and/or vesicular monoamine transporter 2 (VMAT2). More recent work using intersectional genetics and single-cell transcriptomics has shown that many of these neurons also express the machinery necessary to synthesize and/or release additional neurotransmitters (often termed “combinatorial” neurons), in addition to mono-neurotransmitter populations. These multi-neurotransmitter neurons have been identified in the VTA of rodent models, non-human primates, and humans (for a review, see ^1,2^).

Around 30% of adult rat VTA DA neurons express the vesicular glutamate transporter 2 (VGLUT2), a marker of glutamate (GLUT) neurons ^3–6^, and DA/GLUT neurons have been a large focus of research^3,6–11^. While gamma-aminobutyric acid (GABA) synthesizing enzymes GAD1 and GAD2 are largely absent from *Th*+ neurons in the VTA ^12,13^, neurons can release GABA through reuptake (by the GAT1 GABA transporter) and packaging by VMAT or the vesicular GABA transporter (VGAT), introducing a barrier to targeting and understanding the true proportion of DA/GABA neurons ^14,15^. Finally, another subset of neurons express the machinery to synthesize and release DA, GLUT, and GABA ^12,13,16–18^, making up as many as 20% of adult rat VTA *Th*+ neurons ^13^.

In this study, we describe novel cell-type-specific viral tools that enable successful labeling of two previously identified *Th*+ neuronal populations within the VTA. One population is a DA-only neuron, which expresses the mRNA associated with the synthesis and release of DA, but not GLUT or GABA, and is marked by the GTP cyclohydrolase family member, *Gch1*. The other is a DA/GLUT/GABA combinatorial neuron, which expresses mRNA associated with the synthesis and release of DA, GLUT, and GABA, and is marked by an anion transporter, *Slc26a7* ^13,19^. Using these novel tools, we provide the first functional characterization of *Gch1*+ (“DA-only”) and *Slc26a7*+ (“Combinatorial”) neurons, defining their electrophysiological properties, projection targets, and responses to drugs of abuse. These findings help dissociate DA-specific from multi-neurotransmitter signaling and advance our understanding of functional heterogeneity within the VTA.

## RESULTS

### Gch1 *and* Slc26a7 *mark transcriptionally distinct VTA neuronal populations*

Using single nucleus RNA-sequencing (snRNA-seq), our group and others have previously identified two transcriptionally distinct *Th*+ neuron types, a DA-only neuron marked by *Gch1*, and a Combinatorial neuron marked by *Slc26a7* in the rat (**Fig. 1a**; ^2,13,20^) and mouse VTA ^19^. To determine whether these neuron subtypes are reproducibly identified across studies and species, we compared cell clusters identified in independent snRNA-seq datasets generated from rat VTA samples ^13,20^ with those identified in a large (4.3 million nuclei) high-quality brain-wide reference dataset generated as part of the Allen Institute’s Brain Cell Atlas ^21^. Hierarchical comparisons revealed high correspondence between independent rat VTA datasets, with both DA-only and Combinatorial clusters mapping to the “21 MB Dopa” cell class within the Allen Institute’s whole mouse brain taxonomy (**Fig. 1b**). Further comparison identified DA-only neurons map largely onto “0881-0886: SNc−VTA−RAmb Foxa1 Dopa_2-7” supertypes and Combinatorial neurons map largely onto “0880: SNc−VTA−RAmb Foxa1 Dopa_1” and “0887: SNc− VTA−RAmb Foxa1 Dopa_8” supertypes (**Fig. S1a-b**). As in the rat VTA, mouse DA neurons also segregate based on expression of *Gch1* and *Slc26a7* (**Fig. S1c**). The supertypes associated with Combinatorial neurons preferentially reside along the midline of the mouse VTA (**Fig. S1d**), coinciding with previous localization analysis in the rat VTA ^13^. Together, these results indicate that DA-only and Combinatorial clusters represent conserved subpopulations of VTA DA neurons.

**Figure 1.**
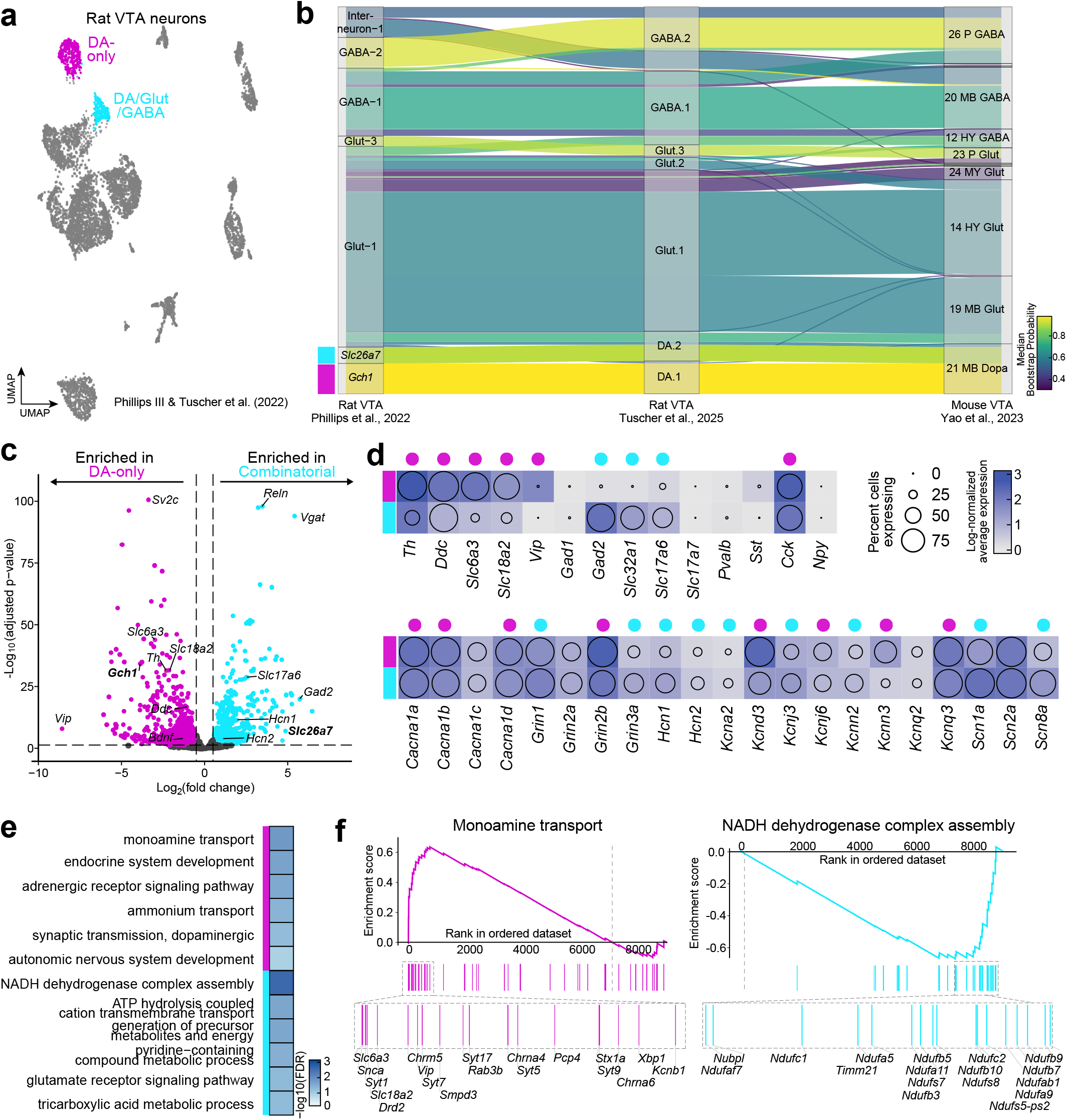
*Gch1* and *Slc26a7* neuronal subpopulations are transcriptionally distinct. **a**, UMAP of rat VTA neuronal populations highlights two distinct clusters previously identified as a DA-only cluster marked by *Gch1* and a DA/GLUT/GABA Combinatorial cluster marked by *Slc26a7*. Adapted from Phillips & Tuscher et al., 2022. **b**, *Gch1* and *Slc26a7* neuronal subpopulations are reproducibly identified between rat sequencing datasets and conserved across rodents. Hierarchical correlation mapping revealed correspondence between rat datasets (Phillips & Tuscher et al., 2022 and Tuscher et al., 2025) and a high-quality mouse reference dataset, with *Gch1* and *Slc26a7* mapping to the “21 MB Dopa” class (Yao et al., 2023). **c**, Volcano plot of DA-only and Combinatorial cluster DEGs reveals extensive transcriptional differences between populations. **d**, Combined heatmap and dotplot analysis shows expression level and percent of cells expressing mRNA associated with neurotransmitter synthesis and release, and selected ion channels associated with action potential and electrophysiologic properties. Magenta and cyan dots above represent genes significantly enhanced in DA-only and Combinatorial populations, respectively. **e**, GSEA analysis shows divergence in enriched gene set pathways. DA-only neurons have an enrichment of genes associated with monoamine transport and DA synthesis, while Combinatorial neurons have an enrichment of genes associated with mitochondrial ATP generation and TCA cycle. **f**, Running enrichment score plot of the highest ranked gene set per population highlights position of specific genes involved in monoamine transport and NADH dehydrogenase complex assembly. Grey vertical dashed line highlights the transition from upregulation (above the x-axis) to down regulation (below the x axis) within the population. A subset of highly enriched genes, alongside their location, are identified below.

Both DA-only and Combinatorial neurons express canonical DA neuron markers relative to other neuronal populations, indicating broadly similar core gene expression patterns. To define transcriptional differences between these populations, we performed differential expression analysis comparing DA-only and Combinatorial neurons in the rat VTA ^13^. Despite the shared DA marker enrichment, differential expression analysis revealed extensive transcriptional differences between DA-only and Combinatorial subpopulations, suggesting broad divergence in their molecular identities and potential function (**Fig. 1c**). This analysis identified 883 genes significantly enriched in the DA-only cluster and 807 genes significantly enriched in the Combinatorial cluster. Examination of canonical neurotransmitter markers revealed distinct molecular identities. DA-only neurons harbored enriched expression of DA markers *Th, Ddc, Slc6a3* (encoding DAT), and *Slc18a2* (encoding VMAT2) compared to Combinatorial neurons, suggesting a DA phenotype (**Fig. 1d**). Conversely, Combinatorial neurons exhibit enriched expression of GABA and GLUT markers, such as *Gad2, Slc32a1* (encoding VGAT), and *Slc17a6* (encoding VGLUT2), compared to DA-only neurons. Neither population had enriched expression of genes commonly associated with major GABAergic interneuron subclasses (including *Pvalb, Sst*, and *Npy*), indicating neither population exhibits canonical transcriptional signatures associated with interneurons (**Fig. 1d**).

Given the vast divergence of DA, GLUT, and GABA neurons in intrinsic and active electrical properties and the role of voltage-gated ion channels in shaping neuronal excitability ^22–28^, we next assessed the expression of voltage-gated sodium and potassium channels and synaptic signaling genes between DA-only and Combinatorial neurons (**Fig. 1c**). I^h^ current is an electrophysiological feature commonly used to identify DA neurons and is generated by hyperpolarization-activated cyclic nucleotide-gated (HCN) channel genes. However, *Hcn1* and *Hcn2* are enriched within the Combinatorial cluster compared to the DA-only cluster (**Fig. 1c-d**), underscoring the need to identify novel DA-specific electrophysiological characteristics.

Given the large number of gene expression differences between DA-only and Combinatorial neurons, we next used pathway analysis to identify biologically meaningful gene sets that differed between these neuronal subpopulations. Gene set enrichment analysis (GSEA) revealed an over-representation of genes involved in monoamine synthesis and transmission in the DA-only population, but enrichment of genes involved in mitochondrial activity and ATP generation in the Combinatorial population (**Fig. 1e-f**). Taken together, these results suggest that DA-only and Combinatorial populations are transcriptionally distinct neuronal subtypes, characterized by divergent expression of genes critical for neurotransmitter synthesis and release, ion channel function, and metabolic processes.

### Novel AAVs allow for cell-type-specific access to DA-only and Combinatorial neuron populations

Existing strategies to target combinatorial neuron populations have relied on intersectional and subtractive genetic approaches to achieve cell-type specificity in multi-gene identity populations ^5,29^. These approaches typically depend on Cre-recombinase and are therefore more widely used in mouse models where Cre driver lines are widely available. Additionally, these strategies often require multiple viral vectors or the generation of custom complex plasmids for each cell type ^5,29^. To overcome these limitations, we designed and generated a promoter-driven viral strategy that enables cell-type-specific access from a single AAV, independent of transgenic lines. This approach enables decoupling of cell-type specificity from transgenic availability and provides a scalable framework for targeting and manipulating molecularly defined neuronal populations.

Given unique expression of *Gch1* and *Slc26a7* within the DA-only and Combinatorial neurons, respectively, we used putative promoter sequences for each gene to drive molecular cargo in a cell-specific fashion ^30–32^ (**Fig. 2a**). Specifically, we generated novel AAVs expressing Cre-P2A-mCherry under a *Gch1* promoter and Cre-P2A-eGFP under a *Slc26a7* promoter (**Fig. 2b**). To validate viral specificity, we performed smRNA-FISH on rat brain coronal sections collected 3 weeks following intra-VTA injection of AAV9-Gch1-Cre-P2A-mCherry and AAV9-Slc26a7-Cre-P2A-eGFP, probing for *Gch1* and *mCherry*, or *Slc26a7* and *eGFP* (**Fig. 2c**). Ninety-four percent of *mCherry*+ cells were *Gch1*+ (**Fig. 2d**), and 90.24% of *eGFP*+ cells were *Slc26a7*+ (**Fig. 2e**) demonstrating high selectivity of these novel promoter-driven AAVs. Cross-labeling between viruses was minimal at the protein level, with only 2.6% of fluorescently labeled neurons expressing both mCherry and eGFP (**Fig. 2g-h**). These results establish that these AAVs achieve cell-type-specific access and fluorescent labeling in the rat VTA in a Cre-independent manner.

**Figure 2.**
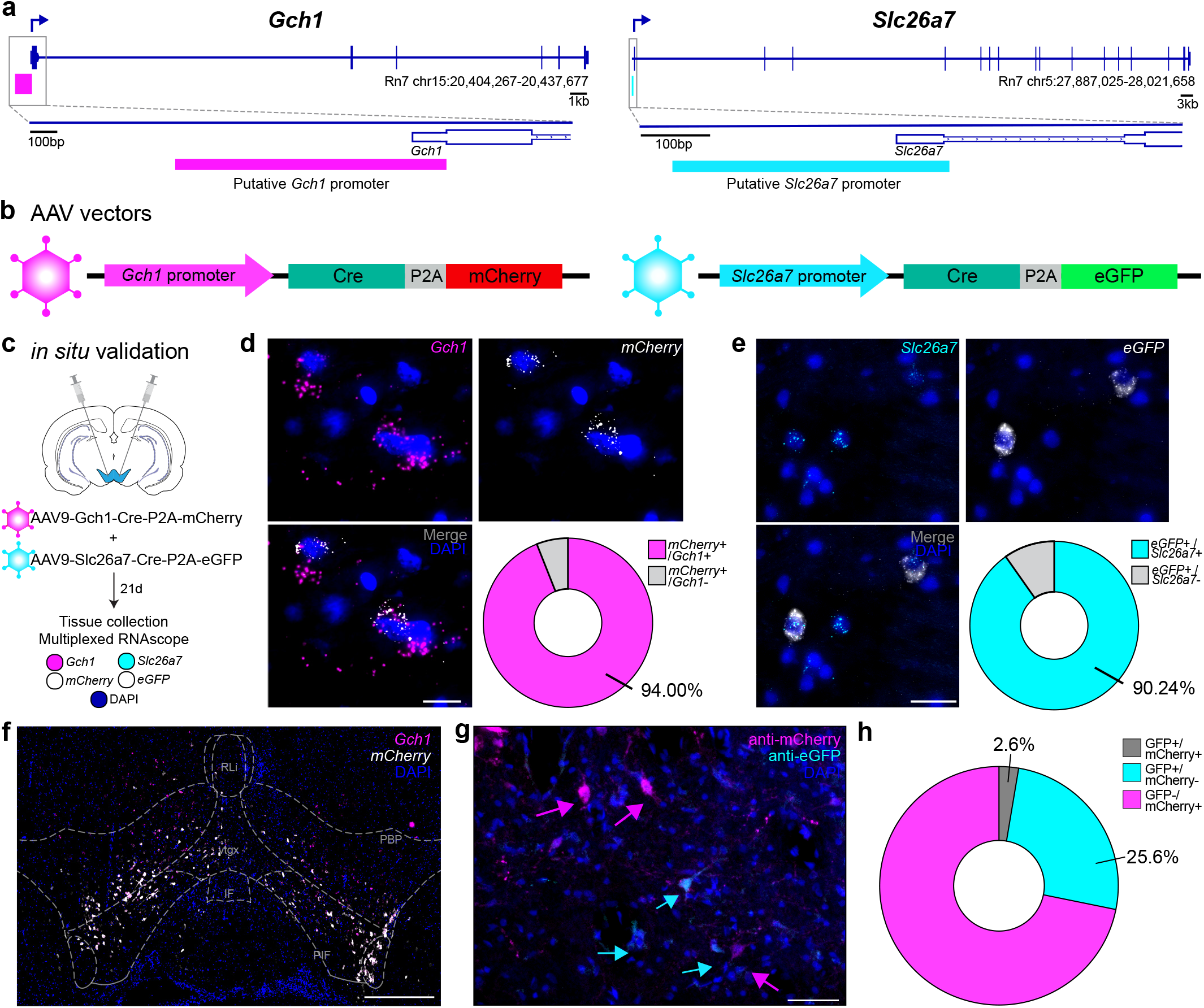
Generation of highly selective AAVs allows for cell-type-specific access of VTA DA neuron subpopulations. **a**, Genomic location of putative *Gch1* and *Slc26a7* promoters. **b**, AAV vectors generated from cell-type-specific promoters transcribe Cre and distinct fluorophores. **c**, Schematic of *in situ* validation of cell-type specificity of viruses. **d**, Representative 100x images of multiplexed RNAscope show *mCherry* transcript puncta are selective for *Gch1*+ cells. Scale bar: 20 µm. **e**, Representative 100x image of multiplexed RNAscope show transcript puncta for *eGFP* localized to *Slc26a7*+ cells. Scale bar: 50 µm. **f**, Representative image of viral spread following intra-VTA injections. Scale bar: 500 µm. **g**, Representative image highlighting mCherry and eGFP protein are in distinct VTA neurons. Scale bar: 50 µm. **h**, Quantification highlights specificity of *Gch1* and *Slc26a7* viruses at the protein level. IF: interfascicular nucleus; PBP: parabrachial pigmented nucleus; PIF: parainterfascicular nucleus; RLi: rostral linear nucleus; vtgx: ventral tegmental decussation

### Combinatorial neurons exhibit higher intrinsic excitability compared to DA-only neurons

Previous studies investigating DAergic electrophysiological properties in the midbrain have relied on classic DA markers, such as TH and DAT, to identify DA neurons. However, given the identification of combinatorial neurons in the VTA that transcribe classic DA markers ^1,13,19^, previous electrophysiological findings attributed to, and used as identifiers for, midbrain DA neurons are now being called into question ^22,33,34^. Sequencing data in the rodent VTA suggests that as many as two-thirds of VTA *Th*+ neurons do not transcribe *Gch1* and may therefore be unable to synthesize DA ^13^. Therefore, previous investigations may attribute electrophysiological results obtained from combinatorial neurons to DA-only neurons, obscuring any differences between these neuronal populations.

To determine if DA-only and Combinatorial neuronal populations differ in basic electrophysiological properties, we performed ex vivo whole-cell patch-clamp electrophysiology in the medial VTA 3 weeks following viral labeling of each neuronal subclass. Our promoter-driven AAVs enabled reliable identification of DA-only and Combinatorial neurons through distinct, non-overlapping, fluorophore expression (**Fig. 3a-b**). Although there were no differences in passive membrane properties (**Fig. 3c**), Combinatorial neurons exhibited a longer latency to fire (**Fig. 3d**) and more pronounced afterhyperpolarization potential (AHP) at rheobase (**Fig. 3e**). Combinatorial neurons further displayed a marked increase in input-dependent excitability and robust sustained firing, evident by greater spike output and the ability to maintain firing at higher current injections. By comparison, DA-only neurons showed reduced ability to sustain firing at elevated current input, and were vulnerable to depolarization block (**Fig. 3f, g**). Following hyperpolarizing current steps (measured at -300 pA), both populations exhibited rebound depolarization (71.43% of DA-only and 40% of Combinatorial neurons; **Fig. 3h**), as evidenced by the presence of at least one action potential, consistent with canonical DA physiology ^35–37^. Together, these data suggest that Combinatorial neurons may have an enhanced capacity for sustained firing under strong excitatory drive, while DA-only neurons may be optimized for more rapid response to inputs but have a limited capacity for sustained depolarization.

**Figure 3.**
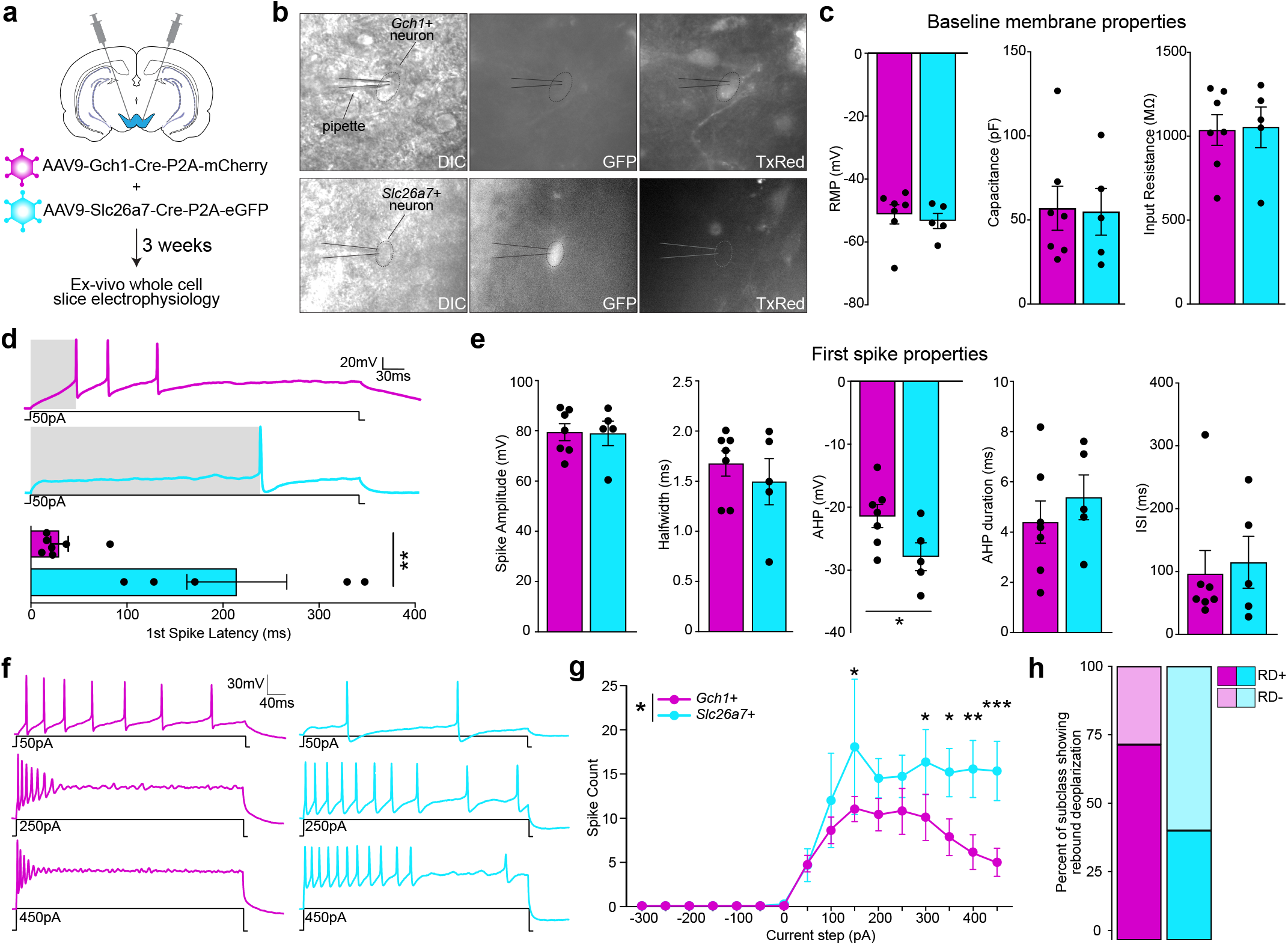
Electrophysiological characterization of DA-only and Combinatorial neurons in the adult rat VTA. **a**, Experimental design. *Ex vivo* whole-cell slice electrophysiology was performed 3 weeks following intra-VTA labeling of DA-only and Combinatorial neurons. **b**, Representative images of patched mCherry+ or eGFP+ neurons within the VTA. **c**, Baseline membrane properties, resting membrane potential, capacitance, and input resistance, did not differ between Gch1 and Slc26a7 neuron populations (DA-only, *n* = 7 cells, 4 animals; Combinatorial, *n* = 5 cells, 4 animals; all *p* > 0.05, two-tailed *t*-test). **d**, Representative traces and quantification of first spike latency at rheobase. Combinatorial neurons had a significantly longer first spike latency at rheobase compared to DA-only neurons (*p* = 0.002, two-tailed *t*-test). **e**, Properties of the first spike differed in AHP potential between DA-only and Combinatorial neurons, but not spike amplitude, halfwidth, AHP duration, or first interspike interval (AHP potential *p* = 0.0490, two-tailed *t*-test). **f**, Representative traces across 3 current steps highlighting the inability of DA-only neurons to maintain firing at high current input. **g**, Input-output curves from whole-cell current clamp recordings reveal significant divergence at high current amplitudes (*p* = 0.0396; step x cell type interaction: *p* = 0.0343, two way ANOVA with Šidák multiple comparisons). **h**, Percent of patched DA-only and Combinatorial neurons exhibiting rebound depolarization (-300 pA). RD+: neuron shows rebound action potential following -300 pA current injection; RD-: neuron does not exhibit rebound action potential following -300 pA current step. \**p* < 0.05, \*\**p* < 0.01, \*\*\**p* < 0.001.

### DA-only and Combinatorial targets differ in projection density

Midbrain DA neurons are known to project to the striatum, with VTA neurons projecting to the nucleus accumbens (NAc) core and shell, where they synapse on GABAergic medium spiny neurons. These NAc projections largely maintain their medial-lateral spatial distribution, such that medial VTA neurons project to more medial NAc, while lateral soma project to more lateral areas of the NAc ^38–40^. VTA projection neurons have also been shown to target the prefrontal cortex (PFC), hippocampus, amygdala, and olfactory tubercle, among other regions ^12,38,41–43^. Through intersectional genetics, previous studies have identified DA/GLUT projections as highly innervating the medial NAc shell ^9,44–46^, while DA/GABA neurons project to the NAc and lateral habenula (LHb) ^16,47^. However, the projection localization of DA/GLUT/GABA combinatorial neurons is unknown. In order to address this gap, and to identify the projection targets of these two subpopulations of DA neurons, we performed immunohistochemistry of virally labeled DA-only and Combinatorial axons (**Fig 4a**).

**Figure 4.**
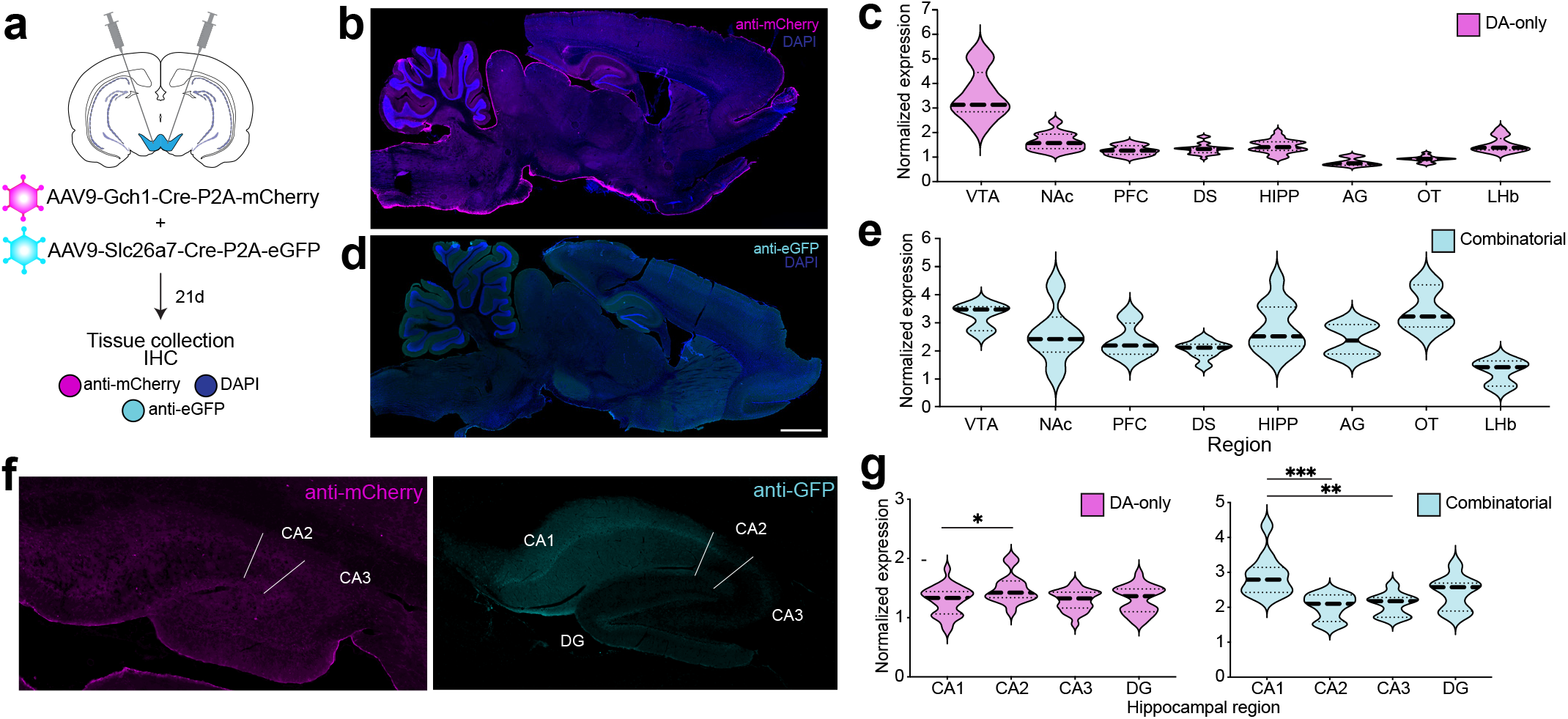
Genetically defined VTA dopamine populations exhibit distinct projection patterns. **a**, Experimental design. Three weeks following intra-VTA viral delivery, perfused tissue was collected and sectioned for IHC. **b**, Representative sagittal section showing DA-only projection regions. **c**, Projection density of mCherry axons normalized to background; large dashed line represents region mean. **d**, Representative sagittal section showing Combinatorial projection regions. Scale bar: 2 mm. **e**, Projection density of eGFP axons normalized to background. **f**, Representative hippocampal projection density images. Scale bar, 1mm. **g**, Projection density of mCherry and eGFP axons within hippocampal regions normalized to background. Projection density of DA-only axons in hippocampal region CA1 differed from CA2 (one-way ANOVA, *p* = 0.0358, with Šídák multiple comparisons), while projection density of Combinatorial axons in hippocampal region CA1 differed from CA2 and CA3 (one-way ANOVA, *p* = 0.0003, with Šídák multiple comparisons). AG: amygdala, DS: dorsal striatum, DG: dentate gyrus, HIPP: hippocampus, LHb: lateral habenula, NAc: nucleus accumbens, OT: olfactory tubercle, PFC: prefrontal cortex. \**p* < 0.05, \*\**p* < 0.01, \*\*\**p* < 0.001.

DA-only neurons exhibited prominent projections to multiple regions implicated in reward processing, with dense axonal labeling within the NAc and PFC. DA-only neurons additionally innervated the hippocampus broadly, with labeled fibers present across CA1, CA3, and DG regions. Beyond corticolimbic targets, DA-only projections were also evident in the LHb (**Fig. 4b-c**). Combinatorial neurons displayed a dense axonal innervation in limbic regions, with prominent projections observed within hippocampal CA1, alongside a high density of axonal labeling within the olfactory tubercle (**Fig. 4d-e**). Interestingly, we found that Combinatorial projections to the hippocampus were significantly biased to CA1, with a higher density of eGFP signal in this region compared to CA2 and CA3 (**Fig. 4f**).

### Combinatorial neurons are uniquely activated by cocaine

Cocaine-induced increases in neuronal firing in VTA DA neurons are dependent on the insertion of calcium-impermeable GluN3A-containing NMDA receptors. Incorporation of GluN3A-containing receptors reduces calcium influx and subsequently decreases activation of small-conductance calcium-dependent potassium (SK2) channels, thereby enhancing neuronal excitability ^48,49^. Although the effects of cocaine on the mesolimbic pathway are well characterized (for a review, see ^50,51^), little is known about how Combinatorial neurons, relative to DA-only neurons, contribute to drug-evoked activity changes.

Given the enrichment of *Grin3a* (encoding GluN3A) and *Kcnn2* (encoding SK2) within Combinatorial neurons (**Fig. 1d**), we hypothesized that this population may be predisposed to heightened activity following acute cocaine exposure. In contrast, another common drug of abuse, fentanyl, engages the mesolimbic circuit through a distinct primary mechanism: disinhibition of DA neurons via µ-opioid receptor-mediated silencing of local GABAergic interneurons ^50–52^. Therefore, we predicted that Combinatorial neurons would be selectively activated following acute cocaine, but not fentanyl, exposure.

In order to test this hypothesis, we assessed drug-evoked neuronal activation using the immediate early gene *Fos* as a functional readout of neuronal activity. To resolve cell-type-specific quantification of stimulus-induced activation, we performed multiplexed RNAscope on coronal sections collected 1 hour following i.p. cocaine (20 mg/kg), fentanyl (0.02 mg/kg), or saline injection, probing for our two marker genes, *Gch1* and *Slc26a7*, and *Fos* (**Fig. 5a**). We found a significant increase in the proportion of *Fos*+ Combinatorial neurons 1 hour following i.p. cocaine (but not fentanyl) injection compared to saline. In contrast, DA-only neurons did not exhibit a significant increase in *Fos* expression following either drug compared to saline (**Fig. 5b-c**). These findings highlight a functional dichotomy between these two *Th*+ VTA subtypes that parallels their transcriptional differences: Combinatorial neurons are selectively engaged by acute cocaine exposure, while DA-only neurons do not show a comparable increase in activity at this timepoint.

**Figure 5.**
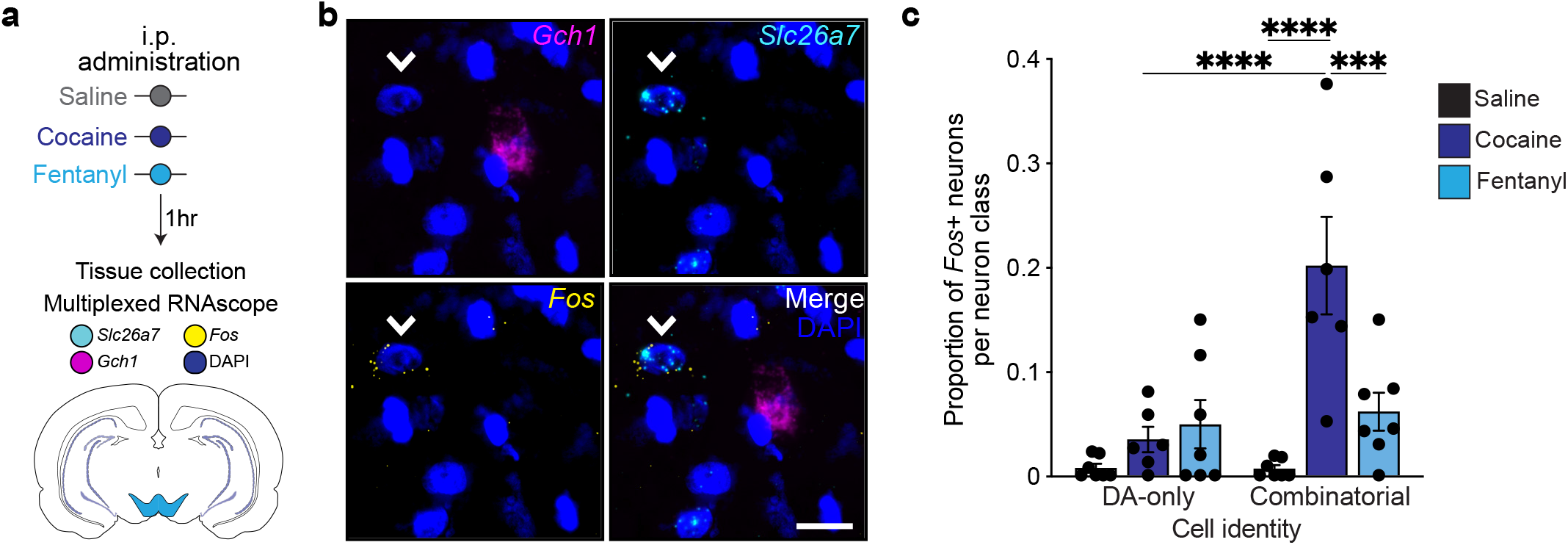
Combinatorial neurons exhibit an increased transcriptional response to cocaine. **a**, Experimental design. **b**, Representative 100x image highlighting a *Fos*+ *Slc26a7*+ neuron and a *Fos*-*Gch1*+ neuron from an animal that received cocaine. Scale bar, 20 µm. **c**, Quantification of the proportion of *Fos*+ neurons within each neuronal class following drug administration (n = 5-7; two-way ANOVA: main effect of drug: *p* < 0.0001; main effect of neuron type: *p* = 0.0142; significant drug x neuronal class interaction: *p* = 0.0006; Šídák multiple comparisons). \**p* < 0.05, \*\**p* < 0.01, \*\*\**p* < 0.001.

## DISCUSSION

In the present study, we further characterize the transcriptional distinction between two *Th*+ neuronal populations within the adult rat VTA, showing strong divergence between DA-only and Combinatorial neurons, with conservation across species. Using multiple orthogonal approaches, including differential expression, pathway-level, and cross-species reference mapping, we show that DA-only and Combinatorial neurons are transcriptionally distinct, rather than variations of a single DA population. In line with previous identification of these populations, DA-only neurons exhibit enhanced expression of DA-associated genes and enrichment of pathways related to DA synthesis and signaling. Conversely, Combinatorial neurons are marked by elevated expression of GABA- and GLUT-associated genes, supporting distinct neurotransmitter identities of these neuron types.

Beyond neurotransmitter phenotype, we further show distinct action potential and ion channel gene expression profiles between the two neuronal populations, providing a transcriptional framework that may underlie the electrophysiological differences observed in this study. In particular, the genes associated with I^h^ current (*Hcn1* and *Hcn2*), often cited as an electrophysiological marker of midbrain DA neurons ^53,54^, were preferentially expressed in the Combinatorial cluster. This finding extends accumulating evidence that I^h^ is an inadequate electrophysiological signature of DA neurons ^22,33,34^, and underscores the need to determine new criteria that distinguishes subpopulations of DA neurons. Finally, the enrichment of genes associated with mitochondrial ATP generation in Combinatorial neurons may explain the electrophysiological and activity-level differences we identified in the present study. Together, these findings build on previous identification of two conserved neuronal populations in the adult rat ^13,20^ and mouse ^19^ VTA. We reveal biologically relevant divergence in neurotransmitter-related and ion channel gene expression supporting the potential for distinct functional roles in motivated behavior.

A major challenge in neuroscience lies in transitioning computational findings to *in vivo* experiments ^55–57^. Here, we used the previously identified highly expressed and selective DA subpopulation markers, *Gch1* and *Slc26a7* ^13,19^, to gain access to each population without relying on Cre-recombinase or intersectional genetics ^5,29^. Utilizing these genetically-driven AAVs, we were able to record from transcriptionally defined neuron classes in the adult rat VTA, enabling direct comparison of electrophysiological properties between *Th*+ neuron types. While DA-only and Combinatorial neurons exhibited similar baseline properties, they differed in key ways, including first spike latency and afterhyperpolarization potential at rheobase and output at high current input, aligning with transcriptional differences in ion channel gene expression (**Fig 1d**). Together, these electrophysiological findings identify a previously unrecognized hyperexcitability phenotype in combinatorial DA neurons.

Consistent with the enhanced mitochondrial ATP generation pathways observed transcriptionally in the Combinatorial neurons (**Fig. 1e-f**), this population was characterized by a tradeoff between sustained firing in elevated energetic demand and delayed recruitment near the threshold for action potential generation. These results refine our understanding of these neuronal subtypes and suggest that DA-only and Combinatorial neurons play distinct yet synergistic roles in the mesolimbic pathway across input levels. DA-only neurons, characterized by rapid responses to lower input but reduced firing persistence at high input, may be optimized for fast, modulatory signaling. In contrast, Combinatorial neurons support sustained output following increased current input. More broadly, these findings highlight the importance of integrating transcriptional identity with physiological characterization to better resolve functionally distinct neuronal subpopulations that have historically been treated as homogenous.

In addition to physiological differences, these two neuronal subpopulations exhibit distinct long-range connectivity patterns. The distinct projection balances observed between DA-only and Combinatorial neurons position these subpopulations to differentially influence key components of motivated behavior. Projections to the NAc and PFC, common to both populations but with an increased proportion of DA-only projections, align with established roles for mesocorticolimbic signaling in salience and action selection (for a review, see ^58,59^). However, the additional weighting toward the LHb in DA-only neurons suggests an enhanced capacity to engage circuits associated with aversive signaling ^17,60– 62^, enabling bidirectional modulation of motivational value. In contrast, the biased targeting of the olfactory tubercle and CA1 from Combinatorial neurons suggests that this population may play a stronger role in contextual and sensory integration, including memory-dependent modulation ^38,63–66^. These differences suggest that the populations are embedded within partially distinct circuit motifs that emphasize valence evaluation or context-dependent motivational control. Because long-range connectivity architecture is considered relatively stable, these data support the interpretation that these populations are organized to engage partially distinct downstream circuits, enabling parallel and specialized contributions to motivated behavior.

We further observed that Combinatorial, but not DA-only, neurons exhibited an increased *Fos* response to acute cocaine exposure relative to saline controls, indicating selective activity-dependent engagement of this population. Cocaine-induced increases in firing of VTA DA neurons depend on the insertion of calcium-impermeable GluN3A-containing NMDA receptors, which reduces activation of SK2 channels and thereby enhances excitability ^48,49^. Although the 1-hour time point examined in the present study is likely too early to reflect de novo transcription and translation followed by receptor insertion, the Combinatorial population exhibits elevated expression of *Grin3a* and *Kcnn2*, which encode GluN3A and SK2, respectively (**Fig 1d**). This enrichment supports the idea that higher basal levels of GluN3A-containing receptors may predispose these neurons to enhanced intrinsic excitability and burst firing following acute cocaine exposure.

Prior studies suggest that combinatorial neurons within the VTA may be preferentially recruited under conditions of heightened stimulus or aversion ^1,18,73^. Although a 20 mg/kg dose of cocaine reliably supports cocaine-paired place preference in rodents ^74^; ^75,76^, a single injection of this dose may also engage aversive or stress-related circuitry. While the present study does not ascertain valence or dose-dependent response, together, the selective activation of Combinatorial neurons may suggest that these neurons contribute to circuit functions extending beyond reward encoding.

These findings align with established cocaine pharmacology in the VTA ^48,49,77,78^ and NAc ^69,70,79,80^, and suggest that DA-only neurons may require downstream circuit engagement, repeated drug exposure, or additional time to drive robust activity-dependent gene expression. In contrast, the selective engagement of Combinatorial neurons points to distinct cocaine-sensitive mechanisms in this population, potentially arising from differences in NMDA receptor subunit composition, intrinsic excitability, metabolic capacity, or afferent input. Together, these results highlight a functional distinction between mono-neurotransmitter DA-only and Combinatorial neurons in their response to drugs of abuse, illustrating how molecular identity can shape drug-evoked cellular adaptations.

Together, these results strengthen the distinction between two previously described VTA DA subpopulations and provide additional characterization of their physiological, projection, and activity-dependent features. The tools generated here establish a foundation for future causal studies to delineate the roles of DA-only and Combinatorial neurons. The transcription of Cre recombinase in a cell-type-specific manner allows for the direct manipulation of these unique neuronal classes through Cre-dependent tools, extending the use of these AAVs beyond cell type identification and labeling. These promoter-driven viral vectors readily bridge the gap between computational identification, physiology, and behavior and can easily be modified to target other unique neuronal populations.

## METHODS

### Animals

All animal model experiments were performed in accordance with protocols approved by the University of Alabama at Birmingham Institutional Animal Care and Use Committee (protocol # 23071 and 20118). Adult male and female Sprague-Dawley rats (35 days old) were obtained from Charles River Laboratories (Wilmington, MA, USA). Rats were co-housed in pairs in plastic-filtered cages with wooden chewing block enrichment in an American Association for Accreditation of Laboratory Animal Care (AAALAC)-approved animal care facility maintained between 22° and 24°C on a 12-hour light/dark cycle with ad libitum access to food (Lab Diet SL3Z Irradiated rat chow) and water. Bedding and enrichment were changed weekly by animal resources program staff. Animals were randomly assigned to experimental groups.

### snRNA-seq data for DA subpopulation comparison

We downloaded the integrated Seurat object from a previous publication of rat snRNA-seq data ^13^ that was aligned to the Ensembl Rn6 genome, and retained all cell type annotations and dimensionality reductions described in the original publication. All downstream analyses here used Seurat v.5.2.1 ^81^ in R v.4.4.0 ^82^. To compare transcriptional signatures across published datasets ^13,20,21^, cluster-specific marker genes were matched based on shared gene symbols, mapping cell types in the current dataset onto cell types defined in Tuscher et al. (2025) using Allen Institute’s Cell Type Mapper, and mapping onto the Allen Institute’s Whole Mouse Brain taxonomy ^21^ using the MapMyCells web interface (RRID:SCR_024671). Cell-type mapping across datasets was visualized using alluvial plots generated in R with ggalluvial v.0.12.5 ^83^.

For pseudobulk differential expression analysis, raw UMI counts were aggregated per sample within each cell type to generate pseudobulk samples. Genes with fewer than 5 total counts across all samples were excluded from testing. Differential expression analysis was performed using DESeq2 v.1.44.0 ^84^. Statistical significance was assessed using a likelihood-ratio test; p-values were adjusted using the Benjamini–Hochberg method, and genes with adjusted p < 0.05 were considered significant. Gene Ontology (Biological Process) enrichment analysis was performed using WebGestalt ^85^ using an over-representation analysis (ORA) approach. The background gene set consisted of all genes tested in the DESeq2 analysis. Enriched GO terms with Benjamini–Hochberg adjusted p < 0.05 were considered significant.

### Cell-type-specific AAV design and production

Adeno-associated viral (AAV) vectors were custom designed by the authors to enable cell-type-specific expression of fluorescent reporters and Cre recombinase. Each construct was based on an AAV-compatible plasmid backbone (pAAV[Exp]-CMV>Cre:P2A:EGFP:WPRE, VectorBuilder) and contained a custom promoter sequence driving expression of a fluorophore (i.e. mCherry or eGFP) and Cre recombinase, as specified in each experiment. Promoter sequences were derived from the rat genome assembly Rn7. Specifically, the promoter region for *Gch1*-targeting virus was cloned from chromosome 15, base pairs 20,437,551 – 20,438,550 (Rn7: Chr15:20437551-20438550) and inserted upstream of the transgene to drive transcription of mCherry and Cre recombinase in *Gch1*+ neuronal populations. The promoter region for *Slc26a7*-targeting virus was cloned from chromosome 5, base pairs 28,021,581 – 28,021,980 (Rn7: Chr5:28021581-28021980) and inserted upstream of the transgene to drive transcription of eGFP and Cre recombinase in *Slc26a7*+ neuronal populations. Genomic coordinates correspond to the Rn7 reference genome.

Final AAV constructs are listed in **Table 1**. Plasmid constructs were assembled and sequence-verified by VectorBuilder (VectorBuilder Inc., Chicago, IL, USA) prior to virus production. Recombinant AAVs were generated by VectorBuilder using triple-transfection in HEK293T cells and purified by cesium chloride gradient ultracentrifugation. Viral genome titers were determined by quantitative PCR and provided by the manufacturer. Final viral preparations were stored at -80ºC until use, and virus was diluted in sterile PBS to the final titer injected on the day of injection.

**Table 1.** AAV viral vectors.

| Virus name | Serotype | Promoter | Genomic coordinates | Transgenes | Titer (GC/mL) | Source |
| --- | --- | --- | --- | --- | --- | --- |
| AAV2/9-Gch1-Cre-P2A-mCherry-oPRE | AAV 2/9 | Gch1 | Rn7:Chr15:20437551-20438550 | Cre, mCherry | $1.11 \times 10^{11}$ | Vector Builder |
| AAV2/9-Slc26a7-Cre-P2A-eGFP-oPRE | AAV 2/9 | Slc26a7 | Rn7:Chr5:28021581-28021980 | Cre, eGFP | $1.11 \times 10^{11}$ | Vector Builder |

### Stereotaxic surgery

Naïve adult male and female Sprague-Dawley rats (Charles River Laboratories) were anesthetized with 4 to 5% isoflurane and placed in a stereotaxic apparatus (Kopf instruments, Tujunca, CA, USA). Anesthesia was maintained at a surgical plane with 2-3% isoflurane, and respiratory rate was monitored throughout surgery and maintained between 35 and 55 breaths per minute. Stereotaxic coordinates were determined using the Paxinos and Watson rat brain atlas (sixth edition), targeting the VTA bilaterally. Under aseptic conditions, burr holes were drilled at anterior-posterior (AP) -5.5 mm and mediolateral (ML) ±2.6 mm, and a 30-gauge stainless steel injection needle (Hamilton Company, Reno, NV, USA) was lowered to a dorsal-ventral (DV) coordinate of −8.41 mm at a 19º angle (all coordinates with respect to bregma). AAV constructs were infused bilaterally using a 10-µL Hamilton syringe driven by a syringe pump (Harvard Apparatus, Holliston, MA, USA), at a rate of 0.25 µL/min, for a total volume of 0.8 µL per hemisphere. Following infusion, the needle was left in place for 10 additional minutes to allow diffusion of the virus before slow retraction. Burr holes were sealed with sterile bone wax, and the incision was closed with interrupted 6-0 monofilament nylon sutures (DemeTECH, Miami Lakes, FL, USA). Postoperatively, rats received buprenorphine (0.05 mg/kg) and carprofen (5 mg/kg) for analgesia, and topical bacitracin was applied to the incision site. Animals were maintained on a heating pad covered with a sterile drape throughout surgery to preserve body temperature.

### smRNA FISH

Animals were briefly anesthetized with inhaled isoflurane and rapidly decapitated using a guillotine (World Precision Instruments, Sarasota, FL, USA). Brains were extracted and submerged in 2-methylbutane over dry ice for 1 min. Flash-frozen brains were removed, wrapped in aluminum foil and parafilm, and stored on dry ice prior to long-term storage at −80°C. Before sectioning, tissue was equilibrated to −20°C for at least 1 hour, then sectioned at −18°C on a Leica CM1860 cryostat (Deer Park, IL, USA) in 10-μm sections.

smRNA-FISH was performed (no. 323136, ACD Bio, Newark, CA, USA) using probes from ACD Bio targeting Rn-*Gch1* (1094971-C4), Rn-*Slc26a7* (1094961-C1), and Rn-*Fos* (403591-C2; for *Fos* quantification), Rn-*Gch1, mCherry* (431201-C3), and Rn-*Slc26a7* (for Gch1-virus validation), or Rn-*Gch1, eGFP* (538851-C2), and Rn-*Slc26a7* (for Slc26a7-virus validation) following the manufacturer’s protocol for fresh-frozen tissue, as previously described ^13,86^. Briefly, tissue sections were fixed in ice-cold 4% paraformaldehyde (w/v) for 15 min, serially dehydrated in ethanol, treated with hydrogen peroxide, and digested with protease IV. Sections were incubated with the appropriate combination of probes (as above) and stored overnight in a 5× saline-sodium citrate (SSC) buffer. Probes were fluorescently labeled with Opal Dyes (Akoya Biosciences, Marlborough, MA, USA; all dyes diluted 1:750). Sections were counterstained with DAPI and coverslipped with ProLong Glass Antifade Mountant (P36984, Thermo Fisher Scientific, Waltham MA, USA).

All images were acquired on a Keyence-BZ800 microscope. Within each experiment, acquisition settings were kept constant for a given objective and channel. Low-magnification (4x) images of the entire section and high-magnification (40x) images of the midbrain were acquired for all animals. Images were stitched using Keyence Image Analyzer software. For regional analyses, rat brain atlas overlays (Paxinos and Watson, sixth edition) were applied to stitched images, and regional crops were generated in Adobe Photoshop.

### smRNA-FISH image analysis

Images were analyzed in ImageJ. DAPI regions of interest (ROIs) were segmented using the StarDist-2D plugin (“Versatile fluorescent nuclei”) mode, probability score of 0.45, and overlap threshold of 0.20. These DAPI ROIs were applied to the other channels (*Gch1, Slc26a7, mCherry, eGFP*, or *Fos*), which were converted to 8-bit images. Mean fluorescence intensity was measured within each ROI, and thresholds for positive signal were determined by identifying the elbow of the mean intensity distribution curve for a given probe, representing the inflection between background and true signal. ROIs exceeding this threshold were considered positive. Data were further processed in R to identify co-expression and classify cell types based on probe overlap.

### Ex-vivo whole-cell patch-clamp electrophysiology

To examine electrophysiologic differences between cell types, whole-cell patch-clamp recordings were conducted as previously described ^23,87^. Fifty- to 70-day-old rats were briefly anesthetized with isoflurane and intracardially perfused with ice cold oxygenated recovery solution (37ºC): 93 mM N-methyl-D-glucamine, 2.5 mM KCl, 1.2 mM NaH2PO4, 30 mM NaHCO3, 20 mM HEPES, 25 mM glucose, 4 mM sodium ascorbate, 2 mM thiourea, 3 mM sodium pyruvate, 10 mM MgSO4(H2O)7, 0.5 mM CaCl2(H2O2), and HCl added until pH was 7.3 to 7.4 with an osmolarity of 300 to 310 mOsmol. Coronal slices (230 μm) were prepared on a vibratome (Leica VT1200S) containing recovery solution and then transferred to a Brain Slice Keeper (Automate Scientific) containing a holding solution at 31 degrees C for at least 1 hour before recording: 92 mM NaCl, 2.5 mM KCl, 1.2 mM NaH2PO4, 30 mM NaHCO3, 20 mM HEPES, 25 mM glucose, 4 mM sodium ascorbate,

1.mM thiourea, 3 mM sodium pyruvate, 2 mM MgSO4(H2O)7, 2 mM CaCl2(H2O2), and 2 M NaOH added until pH reached 7.3 to 7.4 and osmolarity was 300 to 310 mOsmol. Patch pipettes were pulled from 1.5-mm borosilicate glass capillaries (World Precision Instruments; 2- to 5-MW resistance). Pipettes were filled with K-gluconate intracellular solution for intrinsic experiments. K-gluconate composition was 120 mM K-gluconate, 6 mM KCl, 10 mM HEPES, 4 mM adenosine 5*′*-triphosphate–Mg, 0.3 mM guanosine 5*′*-triphosphate–Na, and 0.1 mM EGTA and titrated to a pH of ∼7.2 with KOH. During recordings, slices were transferred to a perfusion chamber and continuously perfused with artificial cerebrospinal fluid at 31ºC at a rate of 4 to 7 ml/min: 119 mM NaCl, 2.5 mM KCl, 1 mM NaH2PO4, 26 mM NaHCO3, 11 mM dextrose, 1.3 mM MgSO4(H2O)7, and mM 2.5 CaCl2(H2O2). All recordings were performed in the medial VTA in mCherry-positive (for Gch1+ neurons) or eGFP-positive cells (for Slc26a7+ neurons) that were visualized using CellSens software (Olympus). Cells were voltage clamped at −50 mV, and 16 current steps were injected starting at −300 pA and increasing in increments of 50 pA per step. Elicited action potentials were recorded in current-clamp mode, counted, and analyzed using pClamp11 (Clampfit, Axon Instruments). All action potential properties were taken from the first generated action potential at rheobase. Series access was monitored throughout recordings to ensure cells were only used if the access had not changed more than 10%. All reagents were obtained from Sigma-Aldrich.

### Immunohistochemistry

Animals were deeply anesthetized with inhaled isoflurane and transcardially perfused with ice-cold 1x PBS (Fisher Scientific) followed by ice-cold fresh 4% paraformaldehyde (w/v, Sigma-Aldrich) in 1x PBS. Brains were removed and postfixed in 4% paraformaldehyde for 24 h, then cryoprotected sequentially in 15% and 30% sucrose (w/v in 1x PBS; Fisher Scientific) until equilibrated as indicated by sinking, and flash frozen in 2-methylbutane over dry ice for 1 min. Flash-frozen brains were then removed, wrapped in aluminum foil and parafilm, and stored at -80ºC until sectioning. Prior to sectioning, tissue was equilibrated to -20ºC for at least 1 hour, then sectioned at −18°C on a Leica CM1860 cryostat (Deer Park, IL, USA) into 30-μm sections. Sections were collected in series across a 12 well plate (Thomas Scientific) containing cryoprotectant (30% sucrose (w/v), 50mM phosphate buffer (Fisher Scientific), and 30% ethylene glycol (Sigma Aldrich) in dH_2_O) as previously described ^88–90^. Sections were stored at -20ºC until use in immunohistochemistry.

Free-floating sections from a single well were rinsed 3-5 times in 1x PBS to remove cryoprotectant, then permeabilized in 0.25% Triton X-100 (v/v in 1x PBS; Fisher Scientific) for 30 min. Endogenous peroxidase activity was quenched by incubating sections in 0.3% H_2_O_2_ (v/v) for 30 min, followed by two rinses in 1x PBS. Sections were blocked in ice-cold blocking buffer for 60 mins on the rocker, then incubated overnight at 4ºC in primary antibody diluted in antibody buffer. The following day, sections were washed twice in 1x PBS + 0.25% Triton X-100 for 5 min each, followed by a single wash in 1x PBS. Finally, sections were incubated in the appropriate secondary antibody (**Table 2**; diluted in antibody buffer) for 2 h at room temperature and mounted with DAPI Prolong Gold Antifade Mountant (P36931; Thermo Fisher Scientific).

**Table 2.** Antibodies and immunohistochemistry reagents.

| Antibody | Vendor and Catalog # | Dilution |
| --- | --- | --- |
| Rabbit anti-mCherry | Abcam Ab183628 (lot: 1047303-25) | 1:1000 |
| Mouse anti-GFP 9F9.F9 | Abcam Ab1218 (lot: 1034873-23) | 1:250 |
| Rabbit anti-GFAP | Invitrogen PA110019 (lot: T12643106) | 1:1000 |
| Goat anti-mouse AF488 | Invitrogen A10667 (lot: 2892441) | 1:350 |
| Goat anti-rabbit AF546 | Invitrogen A11010 (lot: 2902337) | 1:2000 |

| Antibody buffer | Vendor | Dilution |
| --- | --- | --- |
| PBS | Fisher Scientific | 1X |
| Blocker BSA | Thermo Fisher 37525 | 1:10 |
| Goat serum | Thermo Scientific 16210064 | 1:100 |

| Blocking buffer reagent | Vendor | Dilution |
| --- | --- | --- |
| PBS | Thermo Fisher | 1X |
| Blocker BSA | Thermo Fisher 37525 | 1:10 |
| Tween-20 | Fisher Scientific | 1:2000 |
| Glycine | Fisher Scientific | 2.2% w/v |
| Goat serum | Thermo Scientific 16210064 | 1:50 |

All images were acquired on a Keyence-BZ800 microscope. Within each experiment, acquisition settings were kept constant for a given objective and channel. Low-magnification (4x) images of the entire section and high-magnification (20x) images of target regions were acquired for all animals. Images were stitched using Keyence Image Analyzer software. For regional analyses, rat brain atlas overlays (Paxinos, George, and Charles Watson) were applied to stitched images, and mean fluorescence intensity for given ROIs were defined using FIJI and normalized to background, either corpus callosum or other white matter region, for a given image. For a given animal, regional intensity was then normalized to the signal within the VTA.

### Intraperitoneal injections

Naive rats received a homecage intraperitoneal injection of either cocaine (20 mg/kg), fentanyl (0.02mg/kg), or saline. Animals were euthanized 1 hour following injection, and tissue was harvested for smRNA-FISH as described below. All injections took place between 2:00 and 5:30 p.m., and sessions were counterbalanced by sex and treatment condition.

### Drugs

Cocaine hydrochloride (C5776, Sigma-Aldrich, St. Louis, MO, USA) was dissolved in sterile 0.9% sodium chloride and injected intraperitoneally at described doses (20 mg/kg) for smRNA-FISH studies. Cocaine solution was made fresh before injection and was protected from light. Fentanyl citrate (F3886, Sigma-Aldrich, St. Louis, MO, USA) was dissolved in sterile 0.9% sodium chloride and injected intraperitoneally at described doses (0.02 mg/kg) for smRNA-FISH studies. Fentanyl solution was made fresh before injection and was protected from light.

### Statistical analyses

smRNA-FISH expression differences were compared with one-way ANOVA with Tukey’s multiple comparisons test. Electrophysiological differences were compared with unpaired *t* test with Welch’s correction, or two-way ANOVA with Šidák’s post hoc tests where appropriate. Immunohistochemical differences were compared with one- or two-way ANOVA with Šidák’s post hoc tests, or linear mixed-effects model where appropriate. Statistical significance was designated at *α* = 0.05 for all analyses. Statistical and graphical analyses were performed with Prism software (GraphPad, La Jolla, CA), or R (linear mixed-effects model).

## DATA AVAILABILITY

All data needed to evaluate the conclusions in the paper are present in the paper and/or the Supplementary Materials.

## ACKNOWLEDGEMENTS

We thank all Day Lab members for assistance and support.

## AUTHOR CONTRIBUTIONS

Conceptualization: NDF, JJD

Methodology: NDF, ETJ, CEN

Investigation: NDF, ETJ, CEN, LGS, KMV

Formal analysis: NDF, ETJ, CEN, JJD

Software:

Resources: NDF, JJD

Funding acquisition: NDF, JJD

Data curation:

Visualization: NDF, CEN, JJD

Validation: NDF

Project administration: NDF, JJD

Supervision: NDF, JJD

Writing—original draft: NDF, JJD

Writing—review & editing: NDF, ETJ, CEN, LGS, KMV, JJD

---

**Figure S1.**
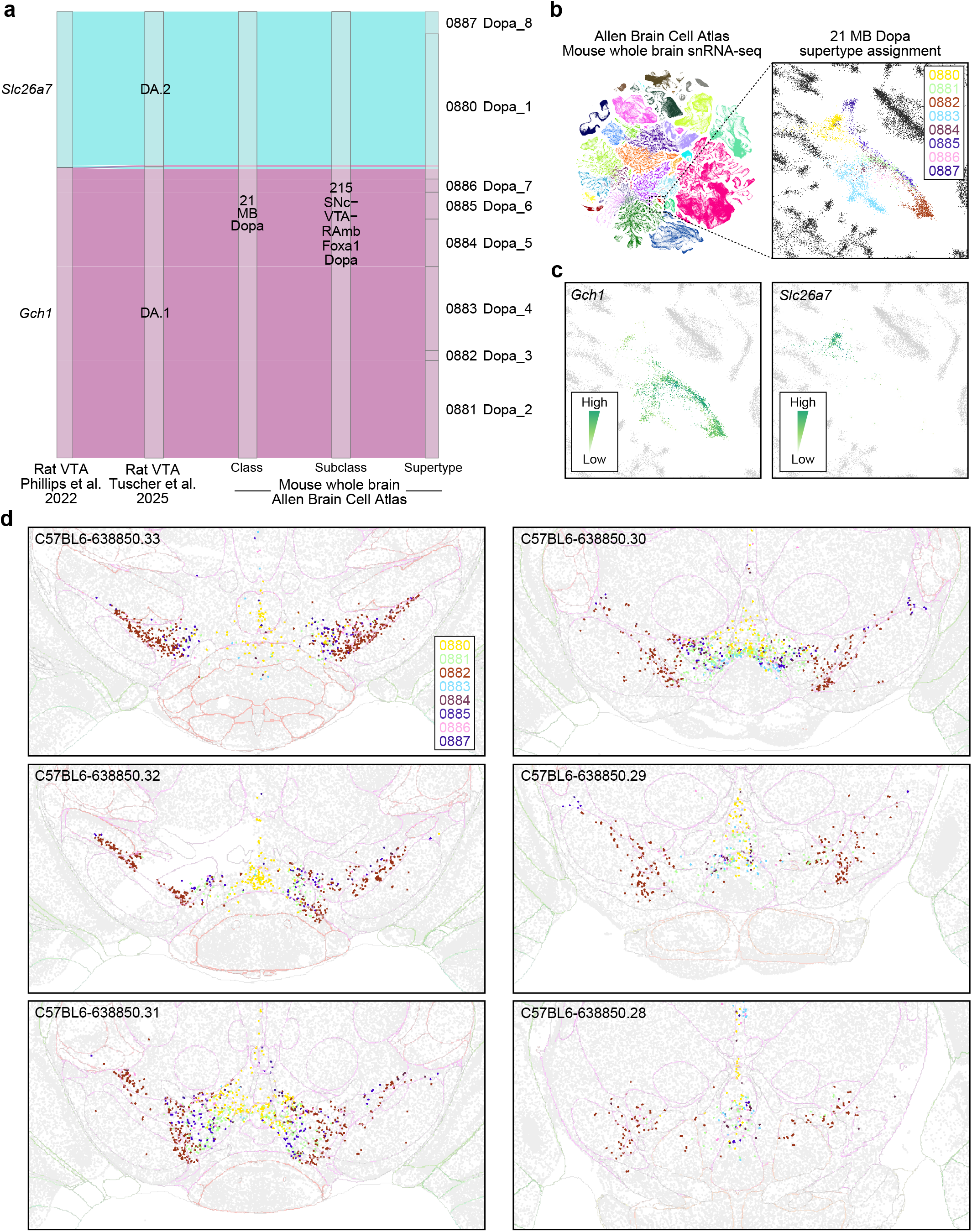
*Gch1* and *Slc26a7* neuronal populations are reproducibly identified between rat snRNA-seq datasets and conserved across rodents. **a**, Alluvial analysis reveals *Gch1* and *Slc26a7* mapping to the “21 MB Dopa” class and “SNc-VTA-RAmbFoxa1 Dopa” subclass (Yao et al., 2023). *Gch1* neurons further map to Supertypes “0881-0886: SNc−VTA−RAmb Foxa1 Dopa_2-7” while *Slc26a7* neurons further map to “0887: SNc-VTA_RAmb Foxa1 Dopa_8” and “0880: SNc-VTA_RAmb Foxa1 Dopa_1” supertypes. **b**, Supertypes of midbrain DA neurons annotated in the Allen Brain Cell Atlas. **c**, Expression of *Gch1* and *Slc26a7* in Allen Brain Cell Atlas. **d**, Location of DA neuron supertypes in the mouse VTA (RRID: SCR_024672).

